# Machine-learning annotation of human splicing branchpoints

**DOI:** 10.1101/094003

**Authors:** Bethany Signal, Brian S Gloss, Marcel E Dinger, Timothy R Mercer

## Abstract

**Background:** The branchpoint element is required for the first lariat-forming reaction in splicing. However due to difficulty in experimentally mapping at a genome-wide scale, current catalogues are incomplete.

**Results:** We have developed a machine-learning algorithm trained with empirical human branchpoint annotations to identify branchpoint elements from primary genome sequence alone. Using this approach, we can accurately locate branchpoints elements in 85% of introns in current gene annotations. Consistent with branchpoints as basal genetic elements, we find our annotation is unbiased towards gene type and expression levels. A major fraction of introns was found to encode multiple branchpoints raising the prospect that mutational redundancy is encoded in key genes. We also confirmed all deleterious branchpoint mutations annotated in clinical variant databases, and further identified thousands of clinical and common genetic variants with similar predicted effects.

**Conclusions:** We propose the broad annotation of branchpoints constitutes a valuable resource for further investigations into the genetic encoding of splicing patterns, and interpreting the impact of common- and disease-causing human genetic variation on gene splicing.

## BACKGROUND

The majority of human genes are spliced, forming a mature mRNA following intron removal and subsequent exon ligation in the spliceosome complex [1]. During this reaction, U1 snRNP and SF1 bind the 5′ splice site (5′ SS) and the branchpoint respectively, and a trans-esterification reaction forms an intron lariat intermediate. A subsequent trans-esterification reaction between the 5′ SS and the 3′ SS removes the intron lariat and forms a spliced RNA product from the flanking exons. Recognition of the sequence-based splicing elements – the 3′ SS, 5′ SS and the branchpoint – by small ribonucleoprotein particles (snRNPs) is a critical step in defining exon boundaries, and therefore the mature mRNA product formed [2].

Mapping of branchpoints generally requires sequencing of the intron lariat following cDNA synthesis. The 5′ SS/branchpoint junction can be traversed by reverse transcriptase, forming a cDNA product that when aligned to the genome has a split and inverted alignment of these elements [3]. However, lariats are rapidly de-branched and degraded, and the vast majority of lariats evade detection by this method [4]. To improve annotations, RNA targeted sequencing was employed to experimentally annotate ~60,000 human branchpoints in ~35,000 introns [5]. This genome-wide annotation revealed the presence of a range of sequence motifs that encompass branchpoints, which have been refractory to previous consensus sequence motif analysis.

Machine-learning (ML) approaches are adept at identifying splicing sequence motifs when sufficient training data is accessible [6]. ML can deconvolute highly complex interactions between seemingly unrelated factors, and therefore can produce a highly complex model capable of predicting where such a feature will occur [7]. Using this previous experimental dataset, we have trained a ML algorithm to identify branchpoints throughout the human genome with high sensitivity and specificity. This approach requires only gene annotations, and does not exhibit an expression-dependent bias in RNAseq that would otherwise limit annotations to highly expressed genes. We provide the branchpoint element maps to current human gene annotations, and a range of additional model organisms, as a resource for understanding splicing.

## RESULTS

### Machine-learning model for human branchpoint element detection

We first designed and trained a machine-learning algorithm, termed *branchpointer,* using experimentally-derived gene annotations [5] to predict potential human splicing branchpoints. To constrain the genome search space and limit unbalancing of classes during training and testing of models, we restricted training to 18-44 nt upstream of 3′ SS annotations, where approximately 90% of known human branchpoints are located [Figure 1A].

**Figure 1.**
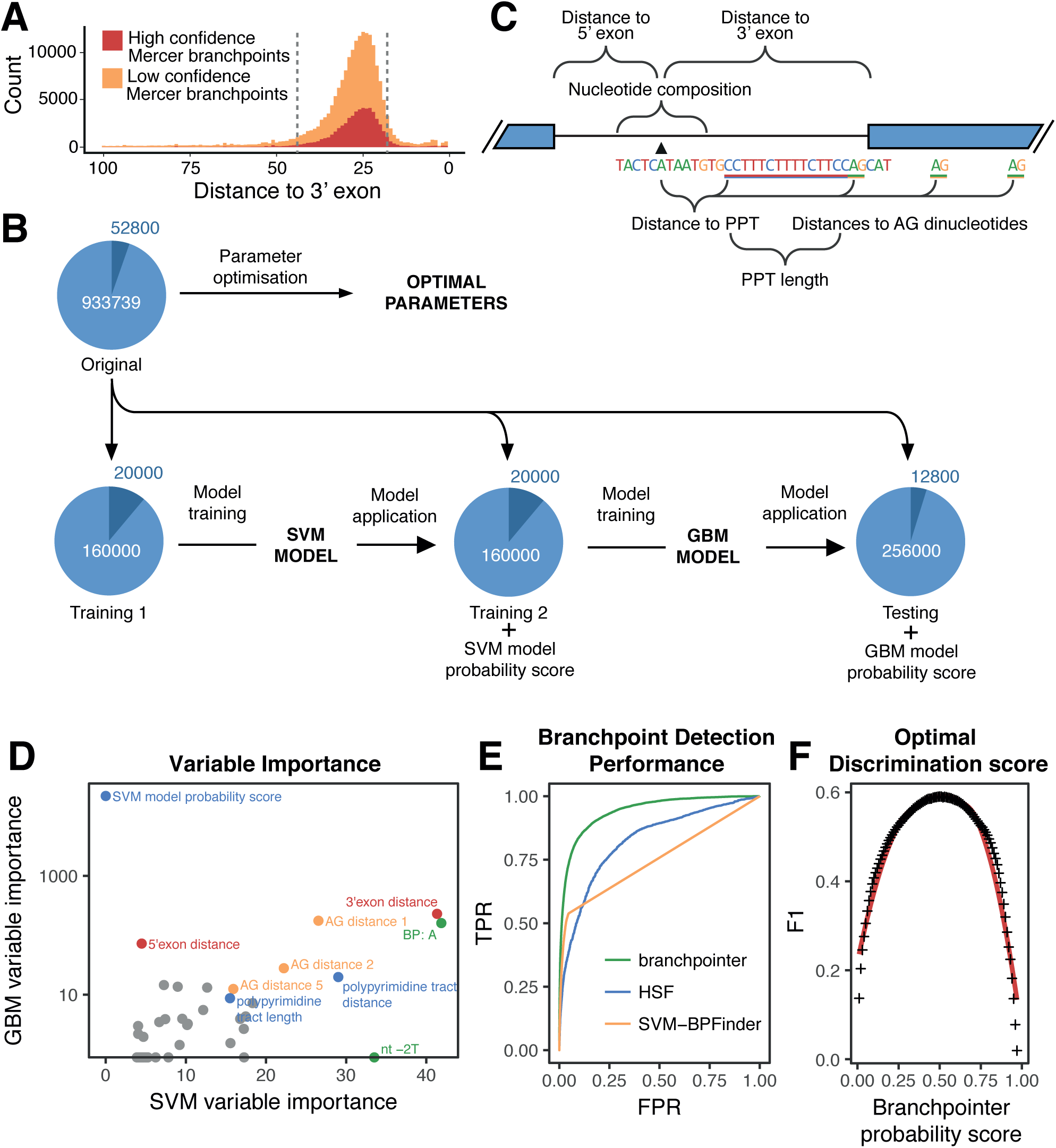
Development and performance of the branchpoint detection model. (A) Branchpoint distances to 3′ exon from the Mercer annotation. Grey dashed lines show the 5th and 95th quantile values for high confidence branchpoints. (B) Schematic of features used in training the branchpoint detection model. (C) Workflow of branchpointer model creation. (D) Importance of variables in the SVM model and GBM model. Only variables remaining after SVM feature selection are included. (E) ROC curves for all predictive methods. TPR: true positive rate; FPR: false positive rate. (F) F1 value for each branchpoint probability score cut-off value.

We generated training and testing datasets using the 52,800 high confidence mapped branchpoints residing within this region as true-positives. To generate a true-negative nucleotide set, we selected intronic regions containing a high confidence branchpoint, and omitted any nucleotide sites with any level of evidence from the experimentally-derived annotations [5]. This selection provided a negative-set of 933,739 nucleotide sites [Figure 1B]. Next, we randomly partitioned 12,800 positive branchpoints and 256,000 negative sites from our dataset to comprise a testing set for subsequent evaluation of model performance. The remaining sites were used to train the *branchpointer* model, a stacked support vector machine (SVM) and a gradient boosting machine (GBM) ensemble.

Previous motifs constructed from a limited amount of data have been used to predict branchpoint sites using a position weight matrix [8]. While this corresponds well to the U2/U12 binding capability of the sequence, other factors are known to influence branchpoint selection including SS definition and the poly-pyrimidine tract (PPT). By incorporating these along with sequence identity [Figure 1C], we were able to rank these encoded features by their relative importance to the SVM and GBM models.

The most significant model features comprise the adenine nucleotide at the potential branchpoint site and the uracil at −2nt relative to the branchpoint [Figure 1D]. These two invariable nucleotides are unique in not undergoing wobble-base pair interactions with the U2 snRNA [5]. Distances to other cis-splicing elements, including 3′ SSs, PPT, annotated 5′exon, and annotated 3′exon were also highly ranked, indicating the relative contribution of gene architecture in branchpoint definition. Notably, features derived from sequence identities at other nucleotides surrounding the tested site – although not as highly ranked – were important for optimisation of overall model performance, supporting evidence that splicing relies on the interplay of *cis* elements to define exon boundaries [2,9].

We assessed the diagnostic performance of *branchpointer* with the testing dataset, returning a receiver operator characteristic (ROC) showing an area under the curve (AUC) of 0.941 [Figure 1E], and a precision-recall AUC of 0.617. Due to an unbalanced number of testing positives and negatives, we additionally assessed *branchpointer* classification performance using the sensitivity, precision, and F1 ratio. Optimal discriminatory class probability for classification was found to be 0.50, with the maximum F1 ratio value of 0.590 [Figure 1F, S1], sensitivity of 0.607, and precision of 0.575. This was found to outperform both currently available methods for branchpoint prediction, and sole use of the UN<u>A</u> motif within the branchpoint window [Table 1, Figure 1E, S2]. *Branchpointer* assigns a probability score to each tested site that reflects branchpoint ‘canonicality’ relative to the initial training dataset [Figure 2A]. For example, branchpoints assigned a probability score >0.95 almost exclusively comprise the canonical UN<u>A</u> motif [Figure S3]. Notably, this probability score does not correlate with U2 binding energy, and a large subset of common branchpoint sites have low predicted U2 binding energy [Figure 2A & B].

**Figure 2.**
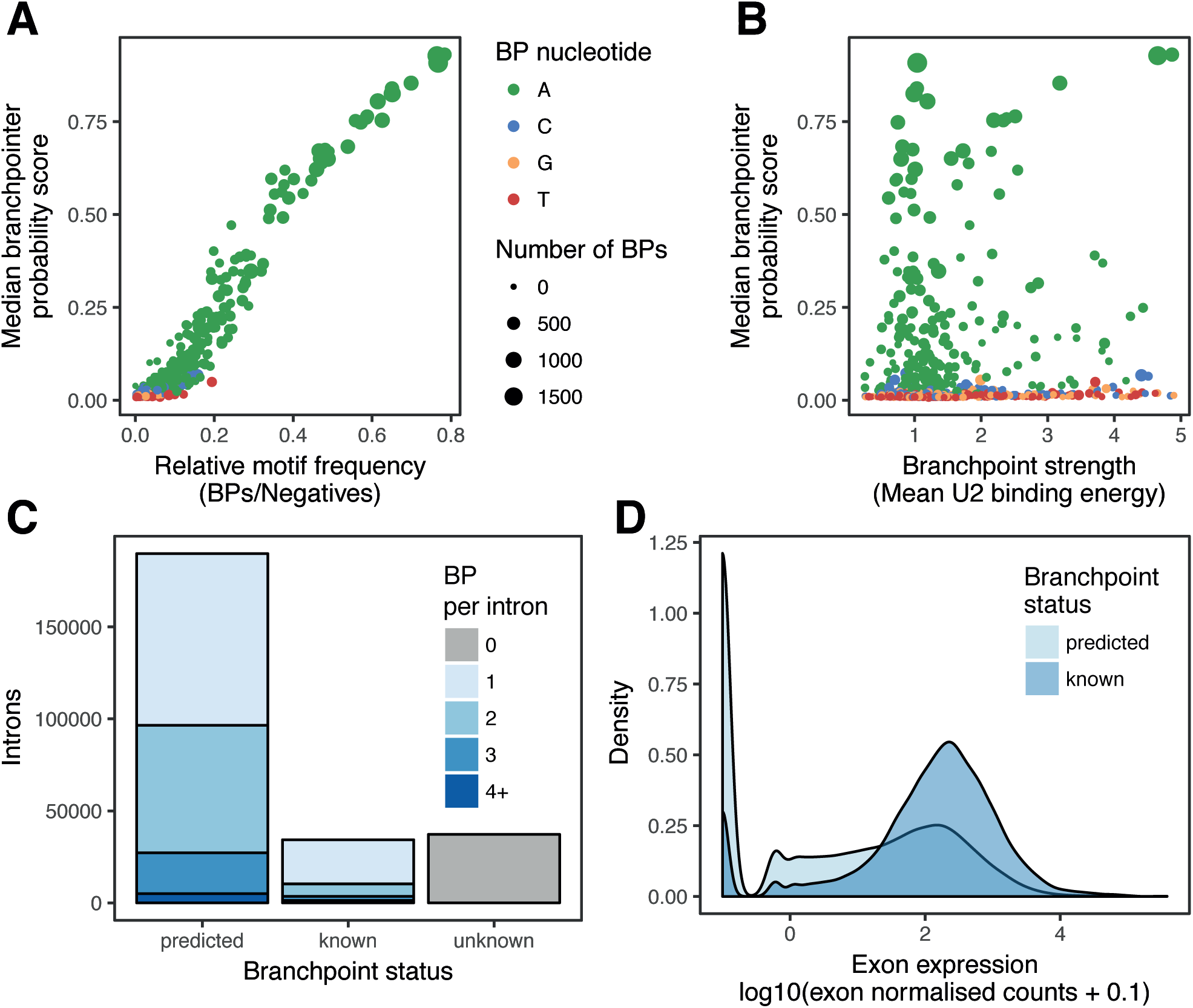
Prediction of splicing branchpoints in GENCODE introns. (A) Median branchpoint probability scores of fivemer motifs by frequency in the training data set. (B) Branchpoint probability scores by branchpoint (BP) strength. (C) Introns with annotated branchpoints from the high confidence Mercer annotation (known), the branchpoint detection model (predicted) and those with no branchpoint (unknown). The default cut-off score of 0.5 was used for predictions. (D) Expression of the associated 3′ exon for introns with known or predicted branchpoints. ENCODE K562 RNA-Seq data was used to generate exon-level counts using STAR and RSEM (see Methods).

**Table 1.** Classification performance of branchpoint predictive methods.

| Method | Accuracy | Sensitivity | Specificity | PPV | NPV | Balanced accuracy | F1 |
| --- | --- | --- | --- | --- | --- | --- | --- |
| HSF | 0.938 | 0.336 | 0.968 | 0.342 | 0.967 | 0.652 | 0.339 |
| SVM-BPFinder | 0.947 | 0.485 | 0.970 | 0.446 | 0.974 | 0.728 | 0.465 |
| *branchpointer* | 0.960 | 0.607 | 0.978 | 0.575 | 0.980 | 0.792 | 0.590 |
| UN<u>A</u> | 0.933 | 0.539 | 0.952 | 0.361 | 0.976 | 0.746 | 0.432 |

PPV: positive predictive value
NPV: negative predictive value

### Branchpoint annotations in the human genome

We applied *branchpointer* to identify branchpoints in all introns in human gene annotations (GENCODEv24 comprehensive human annotation) [10]. Requiring a minimal probability score of 0.5, we are able to annotate an additional 319,143 branchpoints in 189,795 introns [Figure 2C]. This identified a branchpoint element in 85% of tested introns. The remaining introns likely have branchpoints outside the tested branchpoint window; or due to a combination of weak or rare features, are assigned scores lower than our optimal cut-off. Notably, branchpoint annotations are unbiased towards gene type and expression levels, with similar proportions of highly- and lowly-expressed, coding and noncoding genes being annotated [Figure 2D, S4]. In addition to the potential to predict branchpoints in non- or lowly-expressed transcripts, we are also able to predict instances where multiple branchpoints are encoded within an intron, but may not necessarily be utilised [Figure 2C].

### Branchpoints at alternatively spliced exons

We noted that a major fraction (51%) of introns contained more than one branchpoint. These introns are typically shorter than introns with a single branchpoint (Wilcoxon signed-rank, p = 1.48e-163) [Figure S5A]. This trend is also present in the experimentally defined branchpoints, indicating it is not a factor of the window size excluding distal branchpoints [Figure S6], and is observed for both protein coding and long noncoding RNA genes, but not most pseudogene classes [Figure S7]. Introns with multiple branchpoints are also associated with both higher exon expression (Wilcoxon signed-rank, p = 5.00e-08) and intron conservation (Wilcoxon signed-rank, p = 3.82e-21), possibly providing greater mutational redundancy in cases where correct splicing of a transcript is essential [Figure S5B, S5C][11]. Additionally, common variants are less frequent at branchpoints where alternative branchpoints are not encoded within an intron [Figure 3A]. Although there is no significant association with exon skipping (chi-squared, p = 0.13), gene biotype (chi-squared, p = 0.07), or intron biotype (chi-squared, p = 0.09), branchpoint multiplicity is associated with a greater number of annotated alternative 5′ exons [Figure 3B] and a greater diversity of 5′ SS strength [Figure 3C]. This association suggests that multiplicity may also allow more diverse pairing of the 5′ SS and the branchpoint during lariat formation. The strength of splicing elements has previously been correlated with alternative splicing, with strong elements promoting constitutive splicing, and weak elements allowing competition and alternative splicing [12]. Although we were able to observe this correlation at skipped exons for both the 3′ and 5′ SS strength, branchpoint strength at skipped exons is comparable to that of constitutively spliced exons, suggesting strength of branchpoints does not play a major role in exon skipping [Figure S8].

### Human variation at branchpoint sites

Mutations that affect splicing elements can abolish splicing patterns and impact gene expression [13]. To explore the impact of branchpoint mutation on human disease, we searched for ClinVar [14] and GTEx [15,16] SNPs that occurred at annotated branchpoints. As a result, we found five ClinVar SNPs annotated in OMIM [17] that are predicted to delete branchpoints [Table 2]. An additional 248 SNPs had similar predicted effects on branchpoint architecture (see Methods), the majority of which appear to delete branchpoints **[Supplementary Table 2]**. Supporting this prediction, a C-to-T mutation in intron 1 of *Fech* was found to both decrease the strength of the original branchpoint and create a new competing weak branchpoint [Figure 4A-C], and has been previously shown to cause aberrant exon exclusion [18].

**Figure 3.**
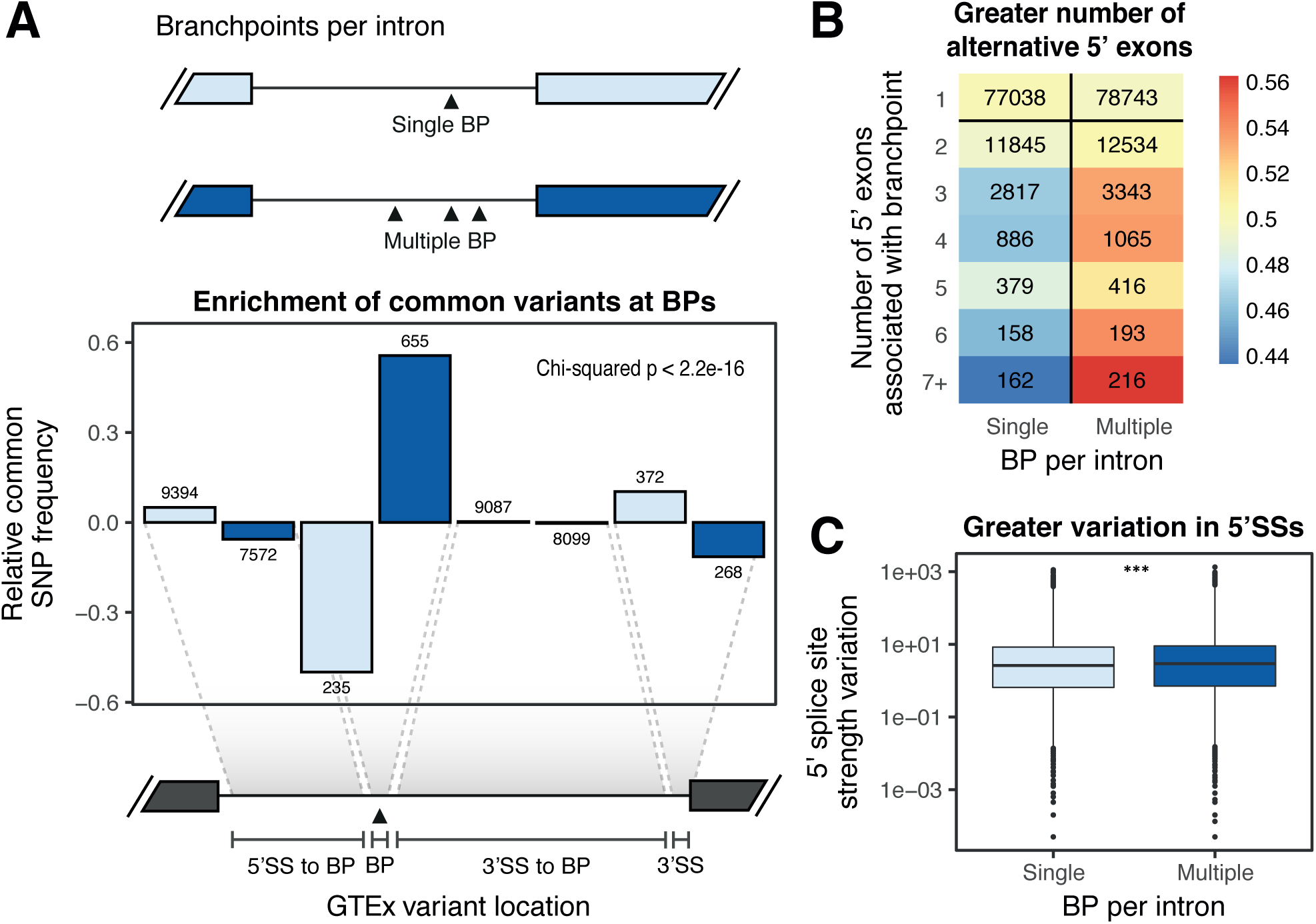
Features of introns with branchpoint multiplicity. (A) Top: schematic and color legend of branchpoint multiplicity within introns. Bottom: Relative frequencies of intronic (1-50 nt from 3′ exon) GTEx SNPs (B) Heatmap of number of alternative 5′ exons annotated in GENCODE transcripts. Color scale is relative normailised percentage. (C) Variation in 5′ splice site strength for introns with multiple annotated 5′ exons. *** p< 1e-06; 3′SS: 3′ splice site; 5′SS: 5′ splice site; BP: branchpoint.

**Figure 4.**
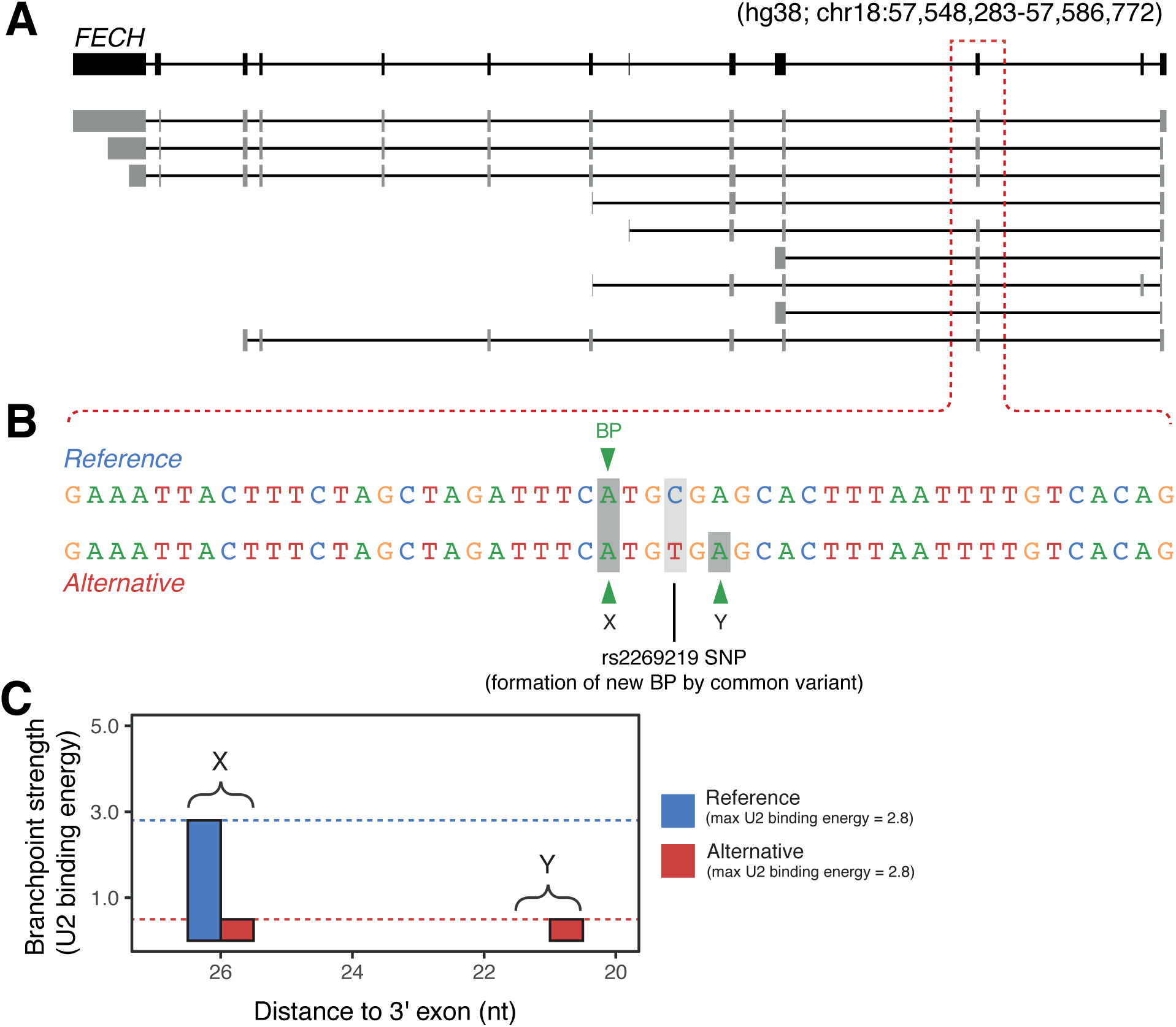
Effect of the SNP rs2269219 on branchpoints in Fech. (A) Gene and isoform structures of Fech either containing or skipping the associated exon (boxed in red). (B) Reference and alternative sequences for the intron containing rs2269219 from −50nt to the 3′ exon. Branchpoints are highlighted in dark grey. Tested SNP is highlighted light grey. (C) U2 binding energy for predicted branchpoints in the reference and alternative sequences (probability score >0.5).

**Table 2.**
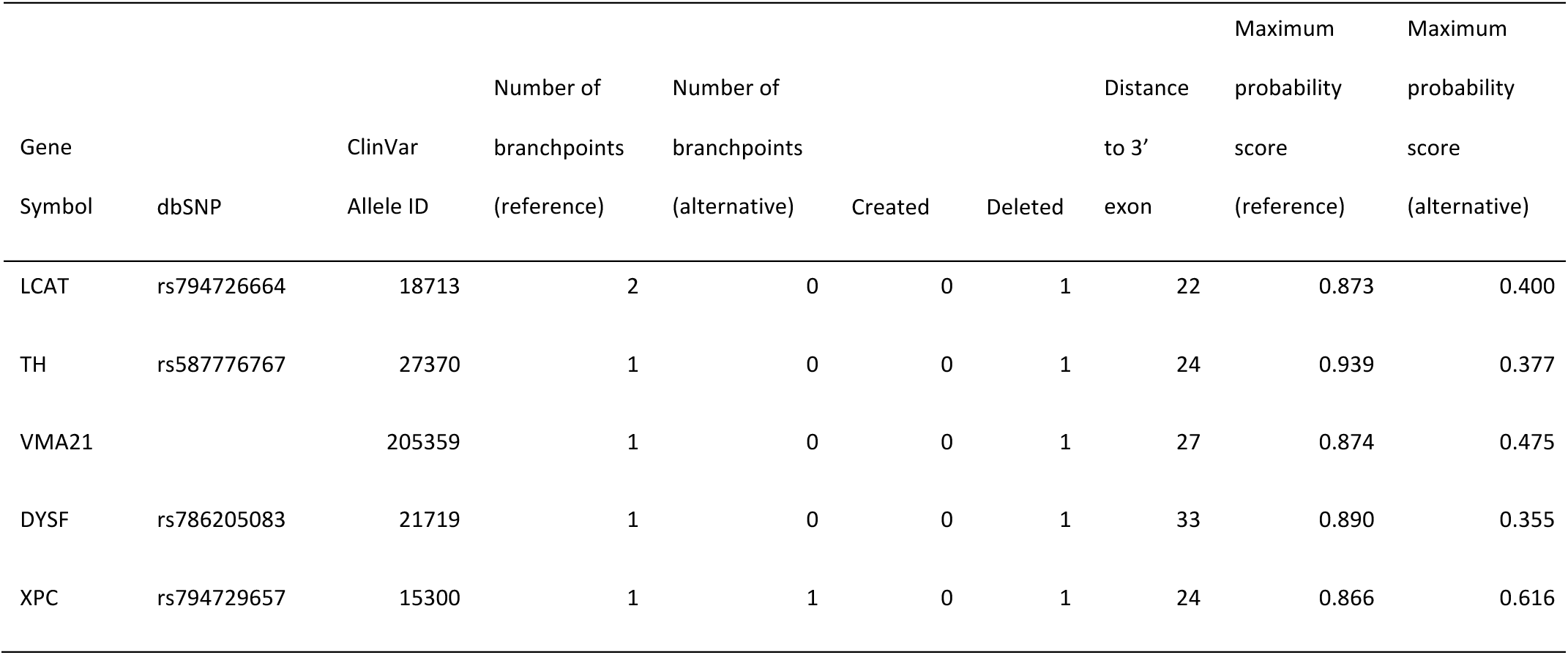
Branchpoint altering SNPs from the OMIM database.

Although there is a relative depletion of branchpoint-affecting common variants, we identified 3870 SNPs as creating and/or deleting a branchpoint **[Supplementary Table 3]**. Unlike disease-associated variants, common variants are less frequent in introns with a single annotated branchpoint than in those with multiple branchpoints [Figure 3A, S9]. Over 20% of branchpoint-altering SNPs in the GTEx catalogue were also associated with an expression quantitative trait loci (eQTL), suggesting that disruption of branchpoint architecture by common (non-disease causing) variants can alter gene expression. This may occur through post-transcriptional regulatory mechanisms like NMD decreasing expression of incorrectly spliced isoforms [19], and may explain the small number of branchpoints (< 2%) associated with splicing quantitative trait loci (sQTLs).

### Branchpoint annotations in model organisms

Given its critical role as a basal element in the splicing reaction, branchpoints elements are conserved in the vertebrate lineage [5]. We also applied the *branchpointer* model to identify branchpoint elements in other model organisms. We used current reference gene annotations to annotate a total 338,322 branchpoints in 194,272 introns in *Mus musculus,* 383,699 branchpoints in 207,847 introns in *Danio rerio,* and 244,149 branchpoints in 133,209 introns in *Gallus gallas,* 258,393 branchpoints in 140,789 introns in *Xenopus tropicalis,* and 84,366 branchpoints in 43,317 introns in *Drosophila melanogaster.* Whilst biased by training in the human transcriptome, these annotations nevertheless are provided as a resource for transcriptome research into model organisms, where branchpoint annotations are largely absent (See ***Availability of data and materials***).

## DISCUSSION

Machine learning is becoming increasingly applied to genomic data, particularly in cases where features are sequence encoded, and thus should be predictable primarily through sequence elements [7]. Splice sites can be readily identified annotated and discovered from exon-exon junction spanning reads from RNA-seq data when the genomic sequence is known [10,20]. This has enabled a sufficiently large catalogue of splice sites to be generated, suitable for training complex ML algorithms [6]. However, annotation of branchpoints currently requires sequencing the intron lariat, and as such, branchpoint catalogues are much smaller. The lack of sufficiently large datasets oversimplified what is now known to be a highly complex and redundant code in humans [5], resulting in multiple biased methods for branchpoint identification [8,21]. Our approach, using the most comprehensive map of experimentally defined branchpoints and ML methodologies is therefore able to capture much more of this complexity, resulting in the most comprehensive and accurate annotation of branchpoints to date.

In the majority of introns, multiple branchpoints are encoded. As branchpoints are an essential element in the splicing of pre-mRNA, a major role of multiplicity may be to provide redundancy in the case of intronic mutations. Multiplicity occurs more often in short, highly expressed, and conserved introns – correlates of genes essential for normal functions in cells [11,22,23]. This may enable faster splicing or mutational redundancy in these important genes. While multiplicity may have a minor role in alternative splicing by promoting alternative 5′ SS usage, we found no significant association with exon skipping events.

Unlike exonic variants, intronic variants can be poorly annotated due to lack of accurate models of intronic sequence elements [24]. We were able to attribute hundreds of clinically associated SNPs with changes in branchpoint architecture. Most of these have not been characterised as affecting branchpoints previously, although all that had and were within the branchpoint window were detected [17]. Disease-associated variants are less frequent and common variants more frequent at branchpoints than splice sites, again indicative that genetic redundancy can act as a buffer to prevent potentially damaging effects of branchpoint deletion.

Genetic variation that impacts RNA splicing can cause disease [25]. However, variations in splicing (sQTLs) are commonly overlooked, as computational tools are typically not sensitive enough to detect most changes in splicing patterns [16,26]. The incorporation of the provided branchpoint annotations in variant identification pipelines provides an additional means to detect potentially damaging variants and inform mechanistic interpretation. With the increasing accessibility to whole genome sequencing in a diagnostic setting there is a growing interest in interpreting noncoding regions of the genome, which are captured but largely discarded in interpretation. Utilisation of accurate branchpoint annotation in clinical interpretation provides a ready opportunity to raise diagnostic yields in whole genome sequencing testing laboratories and sets the stage for future studies to systematically decipher the human genome.

## CONCLUSIONS

We developed a machine-learning model of human splicing branchpoints, which substantially outperforms previous methods for annotation. This model is unbiased to gene type and expression, resulting in the most comprehensive and reliable annotation of branchpoints to date. We found branchpoints are mechanistically distinct from splice sites in terms of their impact on alternative splicing, and multiplicity of branchpoints displays a stronger role in encoding redundancy. Finally, we have made annotations for human and other model species available to the research community (https://osf.io/hrqvq/) [27] to allow future investigations into the biology of this genetic element.

## METHODS

### Dataset generation

Annotated branchpoints from Mercer *et al.* [5] with a single mismatch were used to generate a training set of branchpoints. Introns in Gencode v12 [10] were defined as regions separating two consecutive exons with the same parent gene. Each intron containing an annotated branchpoint was used to generate a positive and negative training set consisting of all locations 18-44 nucleotides downstream of the 3′ exon. Annotated branchpoints contained within this window (52800) comprised the positive dataset, and all other locations not annotated as either a high confidence branchpoint (mismatch at branchpoint) or low confidence branchpoint (split/inverted alignment only) comprised the negative dataset.

### Feature engineering

Nucleotide identity surrounding each location (−5 to +5) was found and converted to a set of dummy variables. Distances to the first 5 canonical AG dinucleotides downstream were calculated. PPTs were defined as strings of sequences starting and ending with a pyrimidine (C/T) and extended until the string consisted of <80% pyrimidines. The PPT for each location was defined as the longest identified PPT between the location and the 3′ exon. Distance to PPT was defined as the distance to from the location to the 3′ end of the PPT. Distance to exon 2 was the distance from the location to the associated 3′ exon. Distance to exon 1 was the distance from the location to the nearest 5′ exon on the same strand.

### Model training and testing

Prior to model training, variables were centred and scaled using the preProcess function from "caret" [28]. SVM models were generated using the "kernlab" package [29], using a "rbfdot" kernel, automatic kpar value and an initial C value of 2. GBM models were generated using the ‘gbm’ method contained within caret. In all cases, classification models were generated which in addition to outputting predicted class could assign a probability to each predicted class. Class probabilities referred to in text are probability of the positive class BP (branchpoint). Probability of the negative class N (not branchpoint) were not used, but is calculated as 1-p(BP). The final stacked model was generated by separating positives and negatives into three subsets – training SVM (20,000BP/120,000N), training GBM (20,000BP/120,000N), and testing (12800BP, 256,000N). First, a SVM was trained using optimized parameters (see below) on the SVM training subset. The SVM model was then applied to the GBM training and the testing subset, outputting BP class probabilities. A GBM model was then trained on the GBM training dataset, with SVM derived BP class probabilities as an additional feature. Performance for both the initial SVM model and the stacked model were evaluated on the testing subset using ROCR [30] and PRROC [31] for ROC and precision-recall curves, and caret::confusionMatrix for classification performance statistics.

The testing dataset was also used to evaluate the performance of *branchpointer* compared to the HSF branchpoint sequence analysis [8], SVM-BP-finder [21] and use of the canonical UN<u>A</u> motif. As HSF assigns scores on motifs alone, all possible eightmer motifs were submitted to the HSF webserver (www.umd.be/HSF3/HSF.html) using “Branch point sequence analysis”. A table of all eightmers and their assigned score is available in Supplementary Table 1. Intronic sequences containing testing sites covering 100nt from the associated 3′ exon were tested using the web server for SVM-BP-finder (http://regulatorygenomics.upf.edu/Software/SVM_BP/). Non-UN<u>A</u> motif containing sites were not assigned a score by SVM-BP-finder, so a dummy score less than the minimum score was assigned. As with *branchpointer,* optimal discriminatory scores were designated as the score that produced the highest F1 ratio for classification of the testing dataset.

### Optimization of training parameters

Performance for each model was initially assessed using a balanced measure of sensitivity and recall - the F1 ratio. Each model was tested on a datasets with a positive to negative ratio of 1:20, which approximates the currently known number of branchpoints within the 18-44nt intron window. Optimal training positive to negative ratios were found by training multiple SVM models on 1000 true positives and varying numbers of true negatives. Optimal C-value for SVM models was found by training models on 5000 positives and 30,000 negatives with varying C values. Feature selection was performed using recursive feature elimination (caret::rfe) with 10-fold cross validation on data subsets containing 2000 positives and 12000 negatives. Following recursive feature elimination variables were sorted in order of importance, and sequentially removed to train a SVM model. Performance of each set of variables was then evaluated on a separate testing subset of the data comprised of 10000 positives and 20,000 negatives. Optimal features were defined as those producing the highest F1 ratio.

### Application to Gencode annotations

Introns within the Gencode annotations (v12, v19 and v24) were defined as outlined in "Dataset Generation". In all introns each nucleotide 18-44nt upstream of the associated 3′ exon was taken and feature values computed. Exon biotype was defined as the biotype of the parent gene. Intron size was defined as the shortest distance between the 3′ exon and an annotated 5′ exon from the same parent gene.

### Model performance

Model performance metrics were evaluated on the testing data subset. We used caret to calculate sensitivity, positive predictive value, accuracy and balanced accuracy of the model as a classifier using cut-off values ranging from 0.01 to 0.99. F1 ratio was calculated manually using the positive predictive value and sensitivity. To compare model performance to SVM-BP-finder, we generated 100nt sequences from all introns within the testing dataset, and submitted these to the SVM-BP-finder webserver for analysis. Only sites contained within the testing dataset were used to compare performance. To compare model performance with HSF, we submitted all possible heptamers to the HSF web tool, and matched these with testing data heptamers. Precision-recall and ROC curves were calculated using PRROC [32].

### Conservation

phyloP 100 way conservation for the hg38 genome was downloaded from UCSC [33,34]. Intron conservation was calculated as the mean value from the 50 nucleotides within an intron up to the 3′ splice site.

### Branchpoint strength

Branchpoint strength was calculated as in Mercer *et al.* [5]. Binding energy of the eightmer surrounding the branchpoint site (−5 to +3) excluding the site to the U2 motif GUGUAGUA was calculated using Rfold from the ViennaRNA suite [35].

### Splice site strength

Splice site strength was calculated using the MaxEntScan score5ss and score3ss tools [36] with the maximum entropy model score as output.

### RNA Sequencing

RNA-seq data from ENCODE K562 cell lines (https://www.encodeproject.org/experiments/ENCSR696YIB/) were trimmed using Trimgalore!, aligned to hg38 using STAR [37] and exon counts from normalised DEXSeq [38].

#### Alternative splicing annotation

Gencodev19 exon annotations were filtered for groups of three sequential exons with either evidence of exon skipping and inclusion (skipped) or strict constitutive splicing (control). Splice junction reads counts were downloaded from GTEx (version 6) [15,16] and used to further refine exon sets for those with evidence of both exon inclusion (>10 junction reads from the middle exon to both the downstream and upstream exon) and skipping (>10 junction reads from upstream to downstream exon), and those with evidence of constitutive splicing only (zero junction reads from upstream to downstream exon and >10 junction reads from the middle exon to both the downstream and upstream exon).

#### Variants

Variants from ClinVar [14] were download on 06/11/2016, and filtered for single nucleotide variants in the GRCh38 assembly. Genotyped and imputed variants were obtained from the GTEx version 6 release (http://gtexportal.org/home/datasets). GTEx splicing QTLs and expression QTLs were obtained from the version 3 and the version 6 data analyses respectively. Variants were filtered for those contained within introns and *branchpointer* was used to assess the effects of SNPs. OMIM [17] SNP variant entries containing "branchpoint" or "branch point", were manually searched for instances where descriptions of the variants referred to either a potential or known effect on a branchpoint. We considered a SNP as potentially altering branchpoint architecture of an intron when at least one branchpoint was created or deleted. Creation and deletion were defined as a change in *branchpointer* probability score greater than 0.2 causing the score to be greater or less than the optimal discriminatory score threshold respectively.

#### Branchpoint prediction in non-human species

Genome sequence and annotation files for *Mus musculus* were downloaded from GENCODE (vM10), and for all other species, annotations were downloaded from Ensembl (Release 85). Branchpoints were predicted in all introns using *branchpointer.*

## LIST OF ABBREVIATIONS

AUC: area under the curve BP: branchpoint
eQTL: expression quantitative trait loci
GBM: gradient boosting machine
ML: machine-learning
NPV: negative predictive value
PPT: poly-pyrimidine tract
PPV: positive predictive value
ROC: receiver operator characteristic
snRNP: small ribonucleoprotein particle
sQTL: splicing quantitative trait loci
SS: splice site
SVM: support vector machine

## DECLARATIONS

### Availability of data and materials

Locations of experimentally defined branchpoints are available as Supplementary Tables from Mercer et al. (2015). Human and mouse gene and genome annotations are available from GENCODE (http://www.gencodegenes.org) and all other species from Ensembl (http://jul2016.archive.ensembl.org). RNA Sequencing data from K562 cells are available from ENCODE (https://www.encodeproject.org) with the accession code ENCSR696YIB. Human genome phyloP conservation scores are available from UCSC at http://hgdownload.cse.ucsc.edu/goldenPath/hg38/phyloP100way/. Splice junction read counts for all tissues, details of GTEx SNPs, sQTLs and eQTLs are available from GTEx (http://gtexportal.org/home/datasets). Clinical variant details are available from ClinVar (https://www.ncbi.nlm.nih.gov/clinvar/). Branchpoint annotations for all species are available in the Open Science Framework repository at https://osf.io/hrqvq/ [27]. Scripts used in the development of the branchpointer algorithm, and scripts used to produce figures are available at github.com/betsig/splice_branchpoints/. The R package *branchpointer* is available at github.com/betsig/branchpointer/. They have also been deposited to Zenodo (https://zenodo.org/) with assigned DOIs: 10.5281/zenodo.192866 (splice_branchpoints) [39], and 10.5281/zenodo.192173 (branchpointer) [40].

### Competing interests

The authors declare that they have no competing interests.

### Funding

The authors would like to thank the following funding sources: Paramor Family Philanthropic funding and Australian National Health and Medical Research Council (NHMRC) Australia Fellowship (1062470 to TRM). BS is supported by an Australian Postgraduate Award. BSG is supported by Cancer Institute NSW Early Career Fellowship 13/ECF/1-45. MED is funded with support from the Kinghorn Foundation. The contents of the published material are solely the responsibility of the administering institution, a participating institution or individual authors and do not reflect the views of NHMRC.

### Authors’ contributions

BS developed the machine-learning algorithm and performed all analyses. BS and TRM conceived the study. BS, BSG, MED, and TRM wrote the manuscript. All authors read and approved the final manuscript.

## Acknowledgements

Not applicable.

---

**Figure S1.**
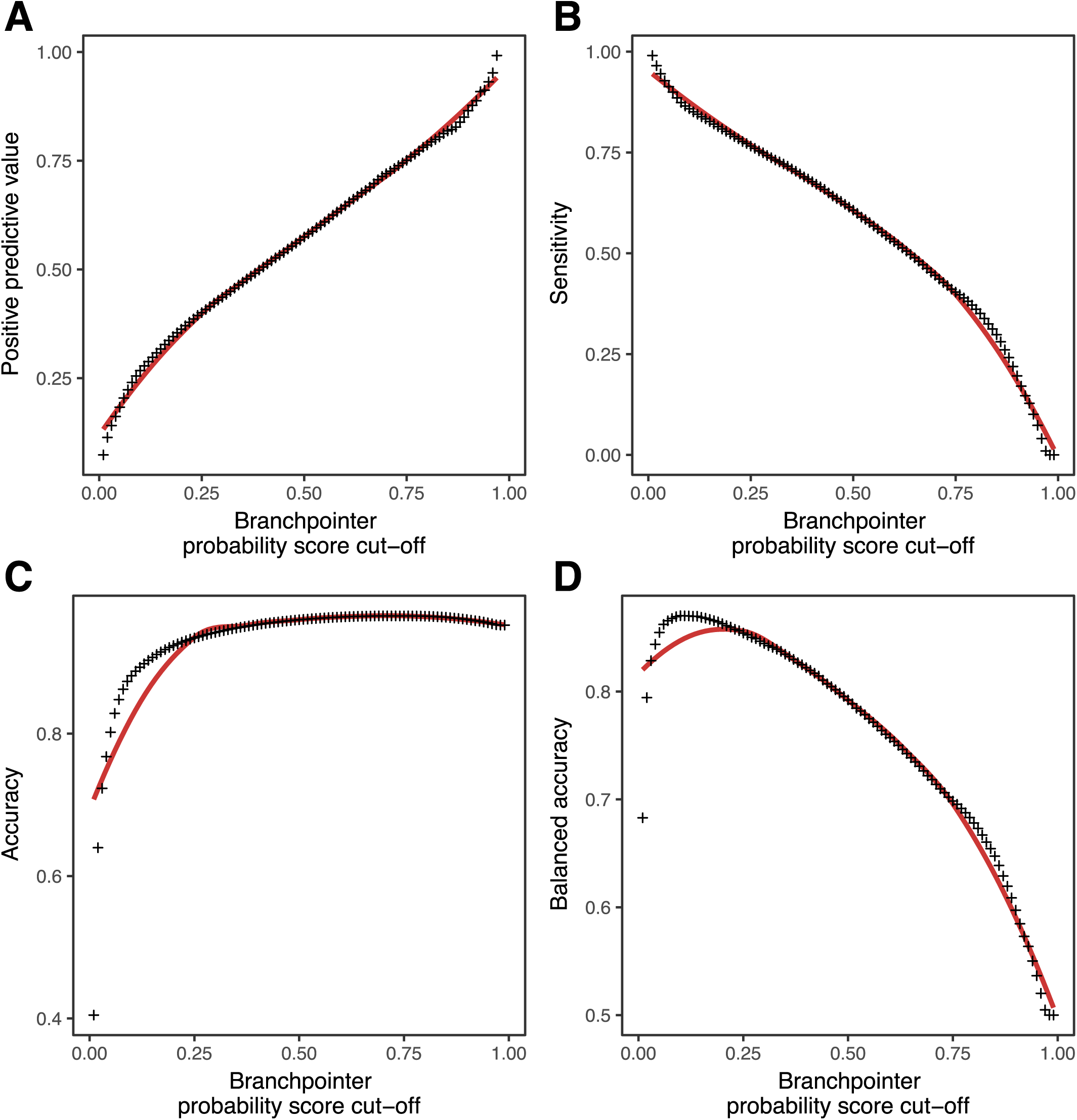
Branchpointer classification performance metrics for probability cutoffs 0.01-0.99. (A) Positive predictive value. (B) Sensitivity. (C) Accuracy. (D) Balanced Accuracy.

**Figure S2.**
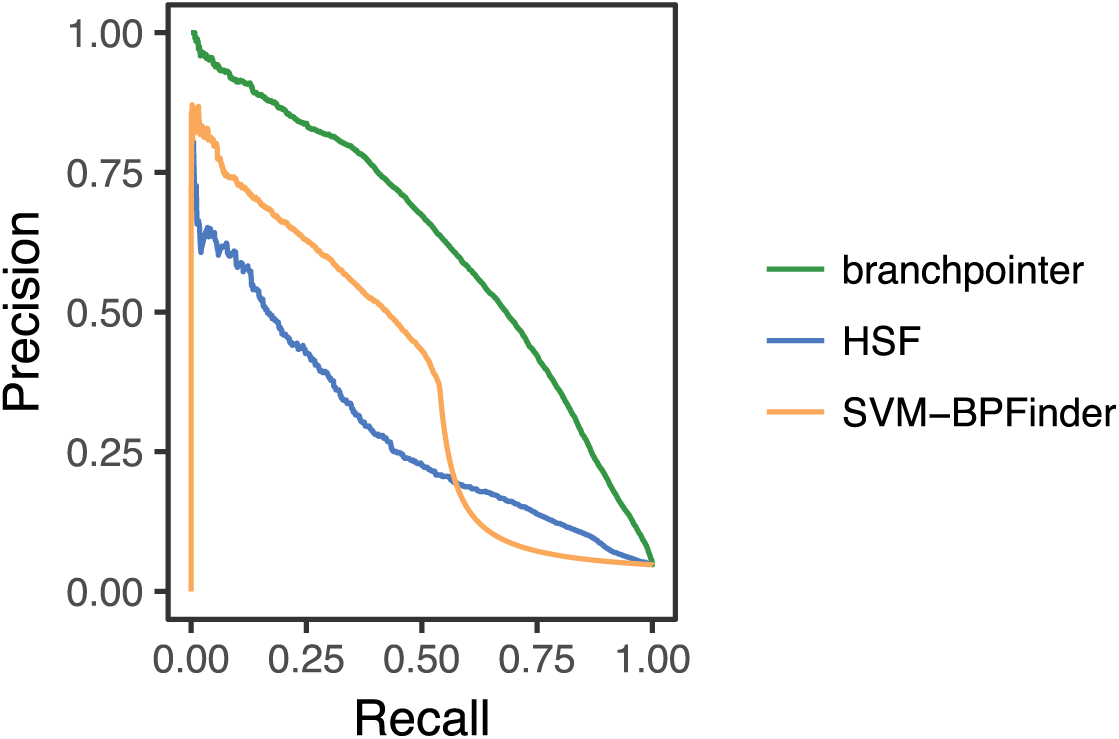
Precision-recall curves for branchpointer, HSF, and SVM-BPFinder.

**Figure S3.**
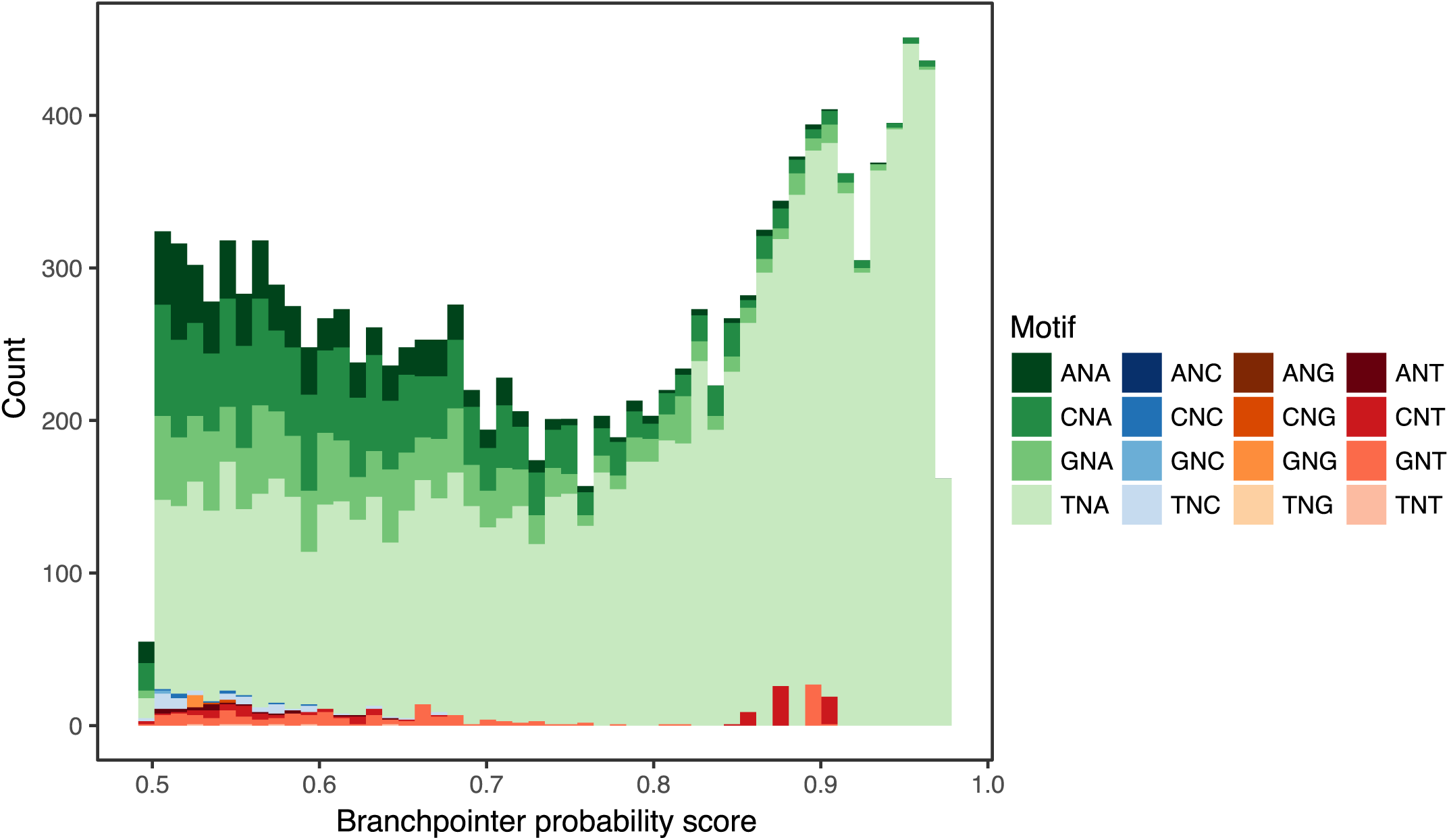
Frequency of each two nucleotide branchpoint motif at each probability score above 0.5. Motif is from position −2 and 0 relative to the branchpoint site.

**Figure S4.**
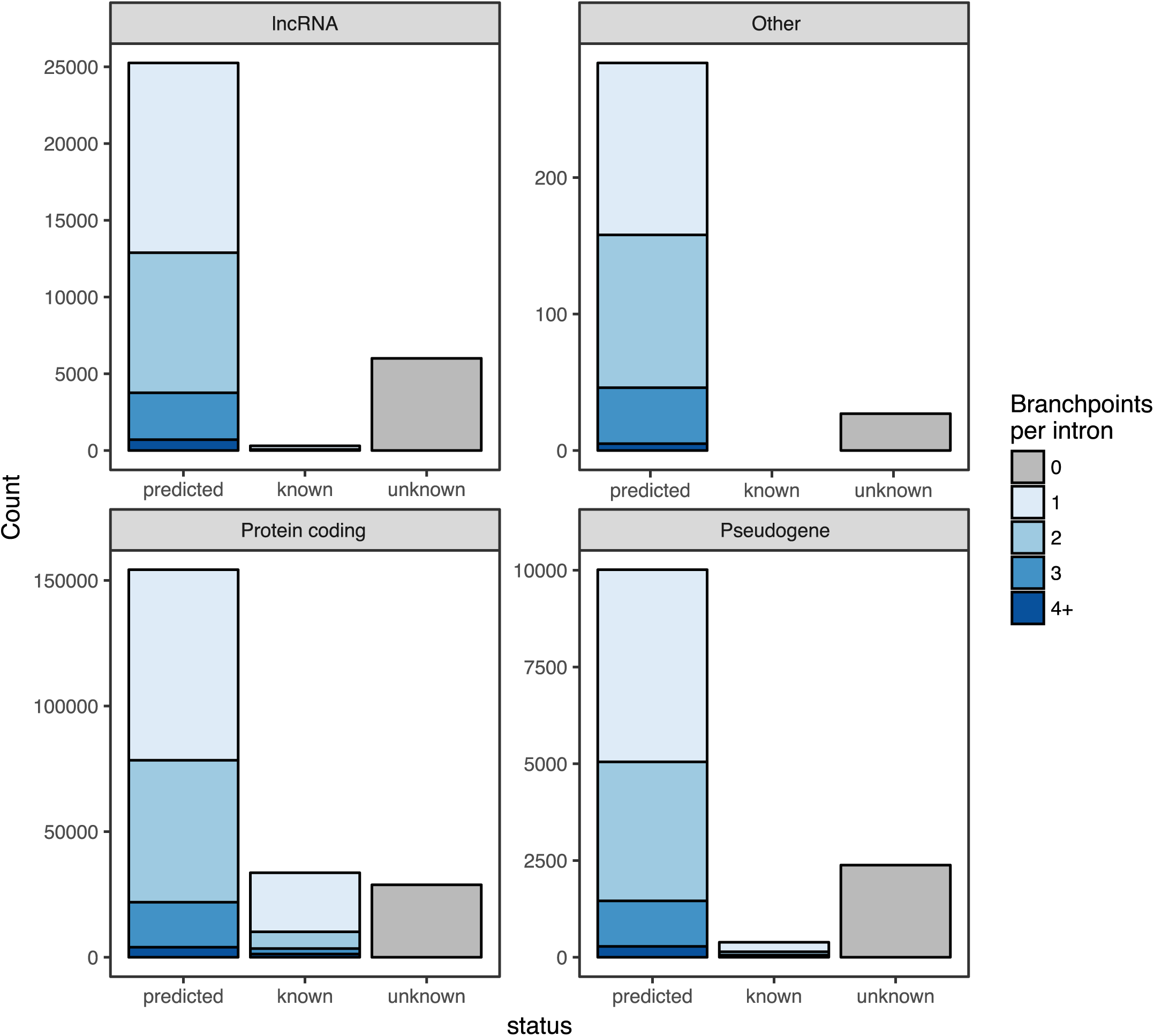
Introns with annotated branchpoints for gene biotypes. Branchpoints are annotated from the Mercer annotation (known), and the branchpoint detection model (predicted) for the gene biotypes long noncoding RNA, protein coding, pseudogenes, and all other biotypes. The default cut-off score of 0.5 was used for predictions.

**Figure S5.**
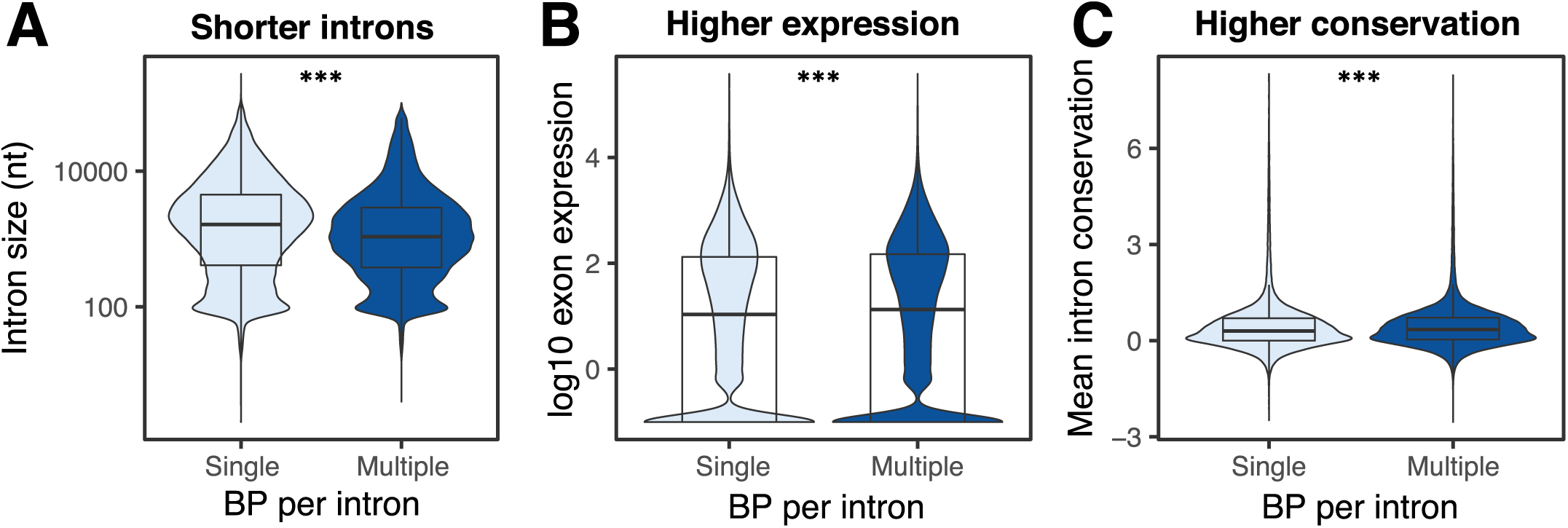
Features of intron usage associated with branchpoint multiplicity. (A) Size of introns with single or multiple annotated branchpoints. (B) Expression of the associated 3′ exon in protein coding genes. (C) Mean intron phyloP conservation. *** p< 1e-06.

**Figure S6.**
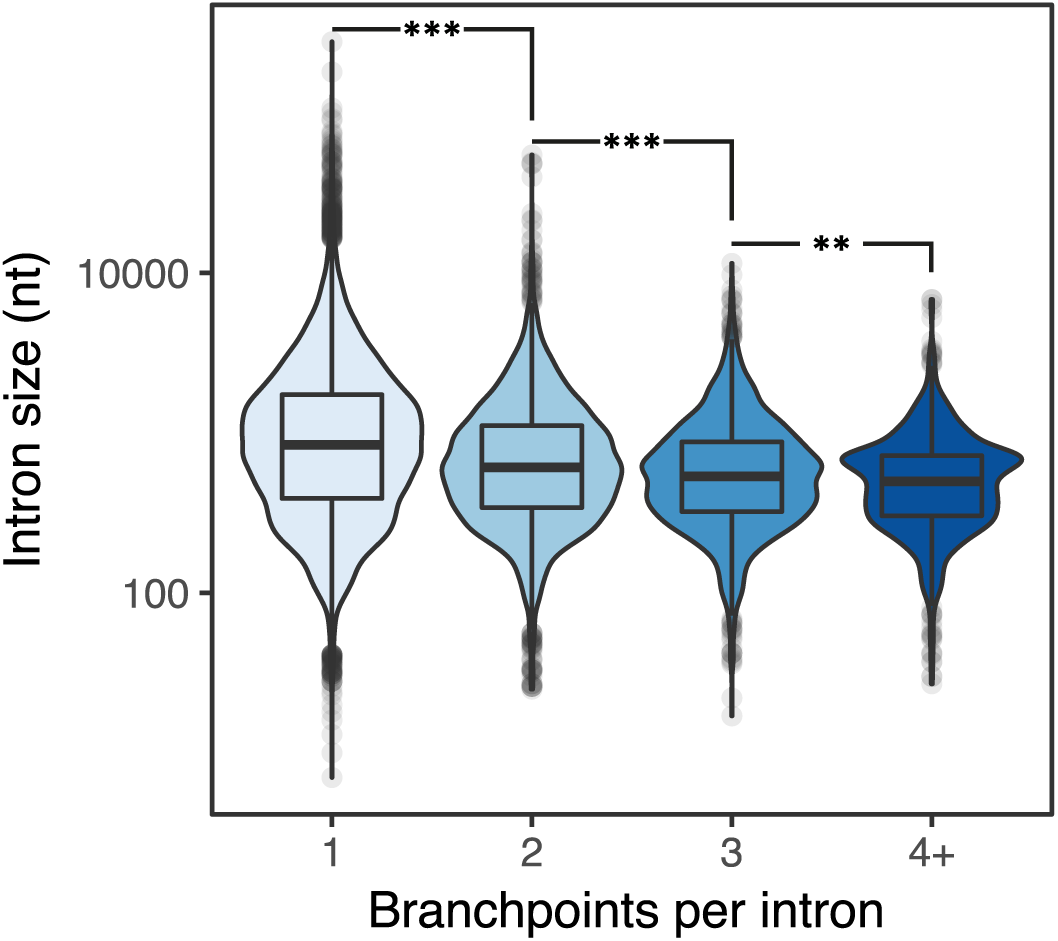
Size of introns with single or multiple annotated branchpoints from the Mercer branchpoint annotation. *** p< 1e-06; ** p< 1e-04.

**Figure S7.**
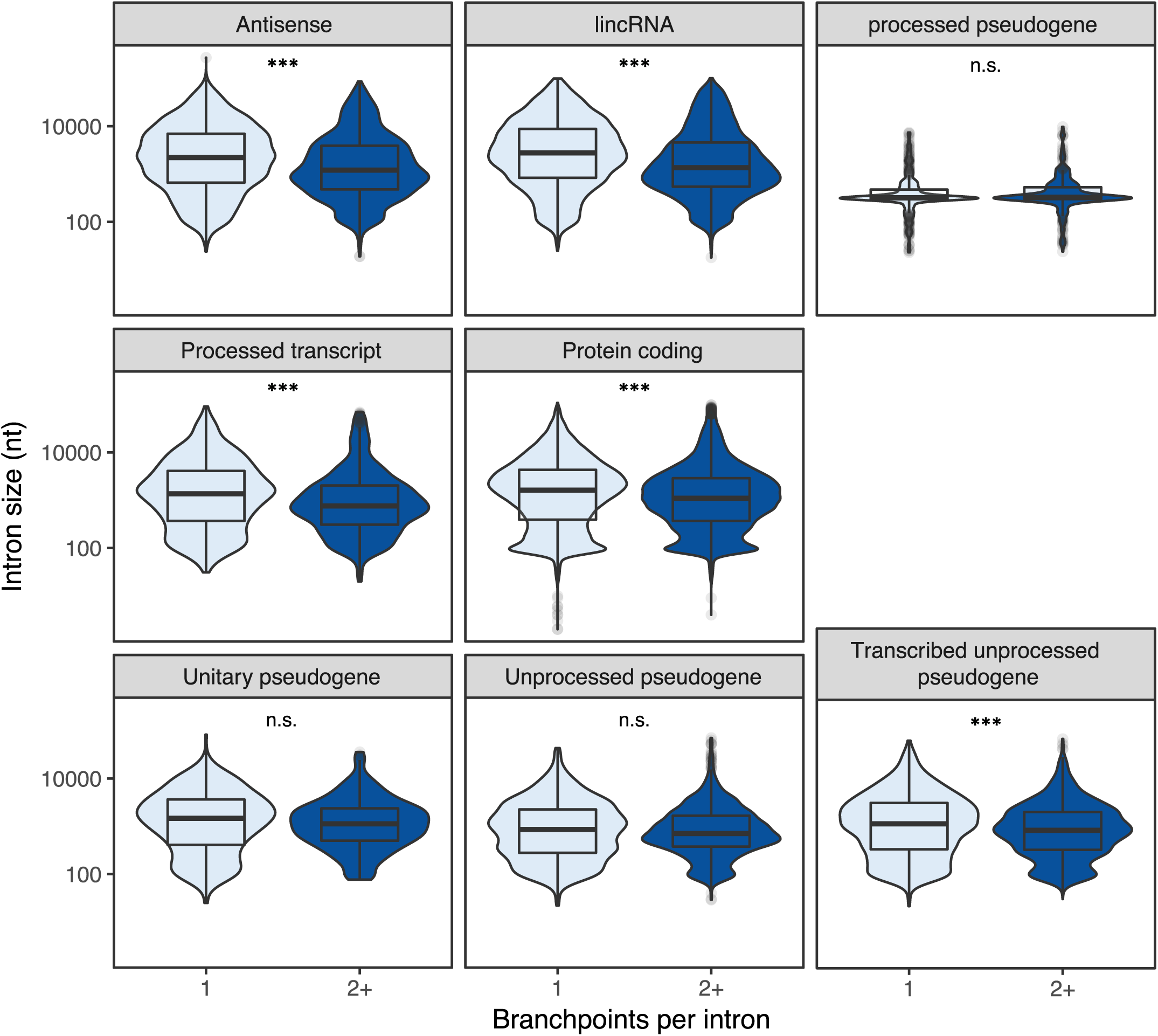
Size of introns with single or multiple annotated branchpoints in parent gene biotypes. Only biotypes with at least 1000 branchpoint-annotated introns are shown. *** p< 1e-06; n.s. p > 0.05.

**Figure S8.**
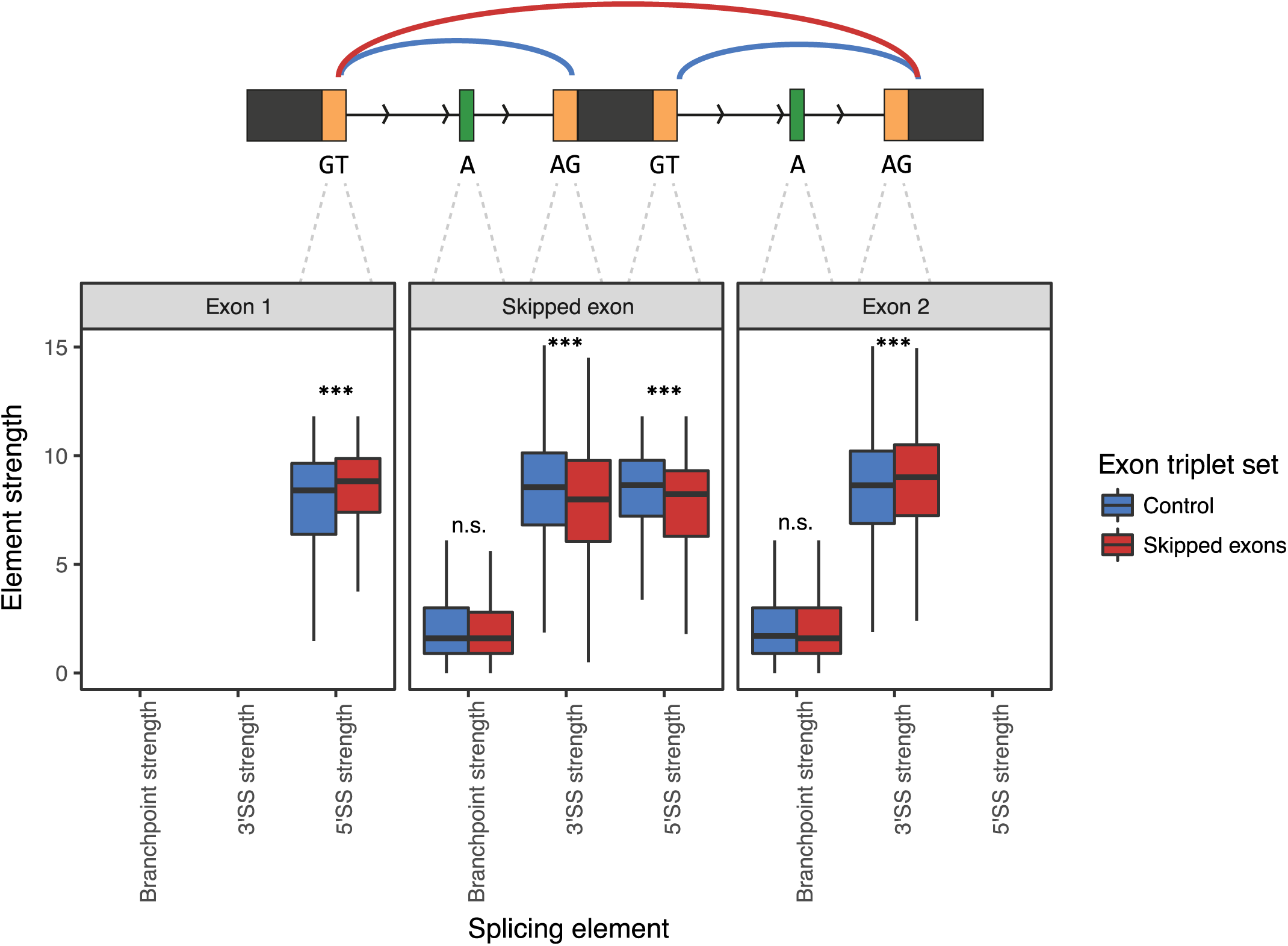
Splicing element strength at constitutively spliced and skipped exon triplets. Branchpoint strength is the maximum U2 binding of branchpoints within the intron. 3′SS and 5′SS strength are calculated using MaxEntScan. *** p< 1e-06; n.s. p > 0.05.

**Figure S9.**
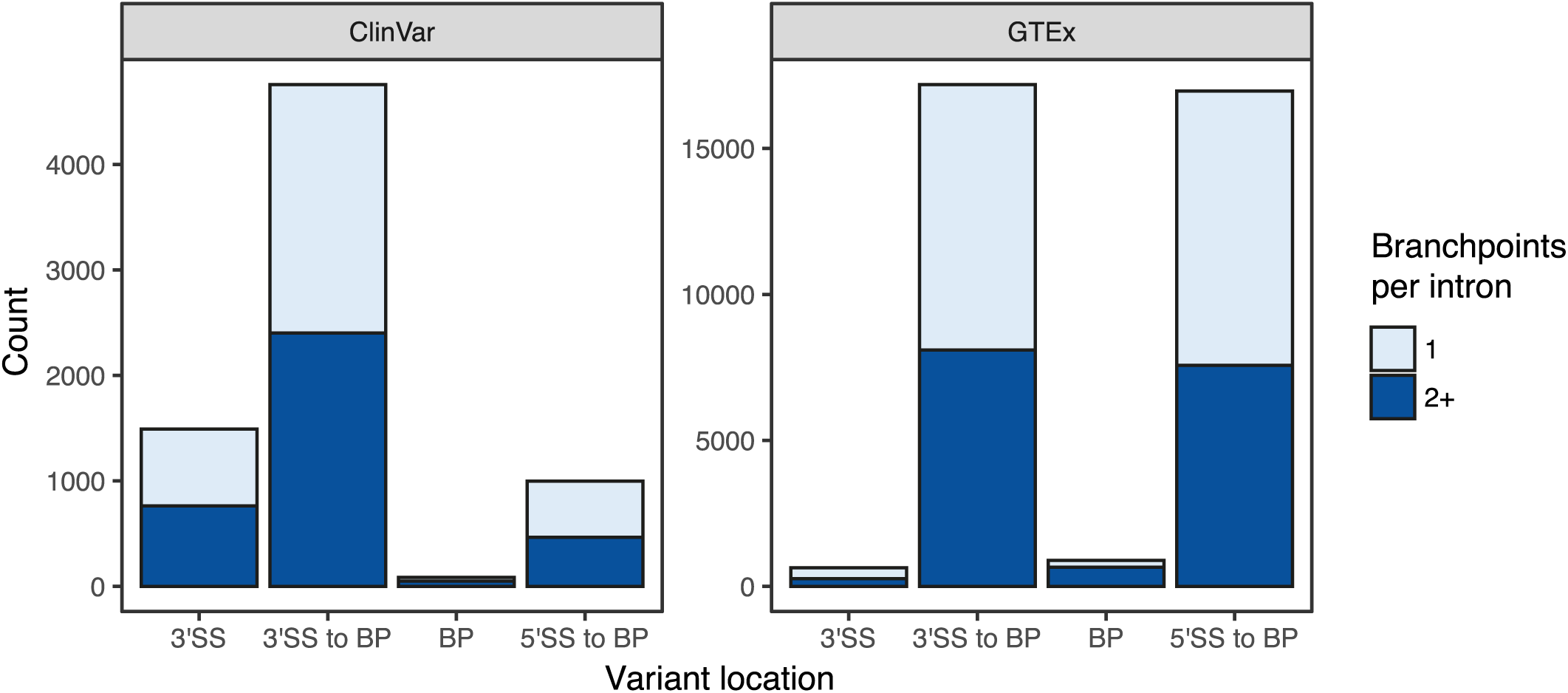
Locations of all intronic (1-50nt from the 3′ exon) ClinVar and GTEx SNPs.

